# T-CLEARE: A Pilot Community-Driven Tissue-Clearing Protocol Repository

**DOI:** 10.1101/2023.03.09.531970

**Authors:** Kurt Weiss, Jan Huisken, Vesselina Bakalov, Michelle Engle, Lauren Gridley, Michelle C. Krzyzanowski, Tom Madden, Deborah Maiese, Justin Waterfield, David Williams, Xin Wu, Carol M. Hamilton, Wayne Huggins

## Abstract

Selecting and implementing a tissue-clearing protocol is challenging. Established more than 100 years ago, tissue clearing is still a rapidly evolving field of research. There are currently many published protocols to choose from, and each performs better or worse across a range of key evaluation factors (e.g., speed, cost, tissue stability, fluorescence quenching). Additionally, tissue-clearing protocols are often optimized for specific experimental contexts, and applying an existing protocol to a new problem can require a lengthy period of adaptation by trial and error. Although the primary literature and review articles provide a useful starting point for optimization, there is growing recognition that many articles do not provide sufficient detail to replicate or reproduce experimental results. To help address this issue, we have developed a novel, freely available repository of tissue-clearing protocols named T-CLEARE (Tissue CLEAring protocol REpository; https://doryworkspace.org/doryviz). T-CLEARE incorporates community responses to an open survey designed to capture details not commonly found in the scientific literature, including modifications to published protocols required for specific use cases and instances when tissue-clearing protocols did not perform well (negative results). The goal of T-CLEARE is to provide a forum for the community to share evaluations and modifications of tissue-clearing protocols for various tissue types and potentially identify best-in-class methods for a given application.

## INTRODUCTION

Tissue clearing refers to methods that increase the physical transparency of biological samples. Through a series of chemical or physical steps, tissue-clearing methods homogenize the refractive index of a sample by removing, replacing, or modifying light-scattering components (e.g., lipids and water). This process increases light transmission and optical imaging depth, making it possible to capture volumetric images of large samples without sectioning the material into thin slices. In recent years, tissue clearing has made it possible to generate high-resolution, three-dimensional (3D) images of entire brains, peripheral organs, bones, and embryos from model organisms (1–18), and large, intact sections of human tissue including brain, breast, prostate, and pancreas (19–22). Data from these images have been used for developmental biology (23), cellular phenotyping (24, 25), identifying and characterizing brain regions governing behavior (26–28), and defining structural aspects of disease pathologies (19, 22, 29, 30).

Selecting and implementing a tissue-clearing protocol can be challenging. Several excellent reviews highlight that no single tissue-clearing protocol (or subset of protocols) is broadly applicable across different experiments. Instead, there are many published tissue-clearing techniques, and each has its strengths and weaknesses (e.g., cost, speed, labeling strategy, toxicity, tissue morphology preservation, and instrument compatibility) (31–40). Thus, the first step for selecting a protocol is to review the literature for similar experiments in similar tissues. Once chosen, the tissue-clearing strategy often must be adapted for specific experimental parameters and research goals (34). Published articles and protocol repositories, such as protocols.io, usually report protocols after they have been optimized and do not provide lessons learned and modifications that were required to generate a successful result.

This article describes the development and implementation of T-CLEARE (Tissue CLEAring protocol REpository; https://doryworkspace.org/doryviz). T-CLEARE is intended to complement publications and protocol repositories and help the community compare and evaluate protocols when designing a tissue-clearing strategy. T-CLEARE is based on the results of a survey that asked researchers to identify tissue-clearing protocols they have used. Respondents were also asked to provide details not usually found in publications or protocol repositories, including protocol effectiveness and ease of use, modifications needed to optimize a protocol for specific circumstances, pre- and postprocessing steps not specific to published protocols, and negative results (i.e., instances when a tissue-clearing protocol did not perform as expected or was not appropriate for the application). T-CLEARE represents each tissue-clearing protocol reported by the community as a flow chart of five steps: fixation, pretreatment/decolorization, labeling, delipidation, and refractive index matching. Each flow chart is included in a tree diagram and organized according to whether it was very successful, successful, or not successful. Users can filter protocols based on whether it was successful, the tissue and organism studied, and the details of five different tissue-clearing steps.

## MATERIALS AND METHODS

### Tissue-Clearing Survey

The tissue-clearing survey was developed by the 10-member BRAIN Initiative 3D Microscopy Working Group (WG) (https://doryworkspace.org/WorkingGroupRoster) between November 2019 and January 2020. The goal of the WG was to establish standards to promote data sharing within the research community. The WG prioritized two approaches for developing standards: (1) to develop standard metadata, which has been described elsewhere (41); and (2) to investigate the feasibility of recommending standard protocols for tissue expansion microscopy. Because of the rapidly evolving nature of the field, the WG agreed that it was premature to recommend one or even a group of tissue-clearing protocols for use by all investigators. In fact, the proliferation of new methods and numerous experimental conditions that must be considered makes it challenging for investigators to select and optimize a tissue-clearing protocol for their own experiments. To address this problem, the WG agreed that a platform was needed to help investigators review and compare protocols. Furthermore, this platform should not be limited to brain studies because the same problems apply to other experiments in other tissues.

WG members reviewed the literature and reached out to colleagues to identify the most widely used tissue-clearing techniques and key experimental parameters that impact selection of a protocol and tissue-clearing success. Based on this input, the subgroup developed a table of commonly used tissue-clearing protocols as a framework for gathering and comparing methods between laboratories. The table included protocol-specific details for five generalized steps commonly used in tissue clearing (34, 42). The table also included a list of experimental constraints that impact selection (e.g., time, cost, use in literature) and success (e.g., specimen, tissue, type of analysis, available equipment).

The tissue-clearing survey was posted on the DORy website (https://doryworkspace.org/tissue_clearing_feedback) for feedback and comment by the larger research community between November 2020 and January 2021. Potential respondents were identified by searching the NIH RePORTER (43) and the BRAIN Initiative website (44) for funded projects that included keywords for relevant microscopy techniques. These investigators were sent an email describing the project and asking them to provide feedback. Respondents were asked to review the table of eight commonly used, published protocols (**Figure 1**). The name of each published protocol was listed in column headers while the five generalized steps were listed in the row headers. Each cell briefly summarizes some of the published details for that protocol for that step. The table also classifies the individual protocols according to mode of action (techniques based on hyperhydrating solutions, tissue transformation, and organic solvents). Respondents reported the tissue-clearing protocol they used by either selecting a cell from a published protocol or specifying modifications using the “other” boxes (**Figure 1**). Respondents were asked to provide a citation (e.g., publication reference, DOI) for any details in the “other” boxes because these represented modifications to one of the protocols listed or use of a protocol not included in the table. In addition, respondents were asked a range of question to capture additional details about their experience (“Was the protocol successful?”, “Why was the protocol chosen?”, “Are you still looking for better tissue-clearing protocols?”) and their experiment (e.g., specimen type, tissue type, feature of interest, label type, type of analysis, level of analysis). A total of 29 respondents provided feedback.

**Fig 1.**
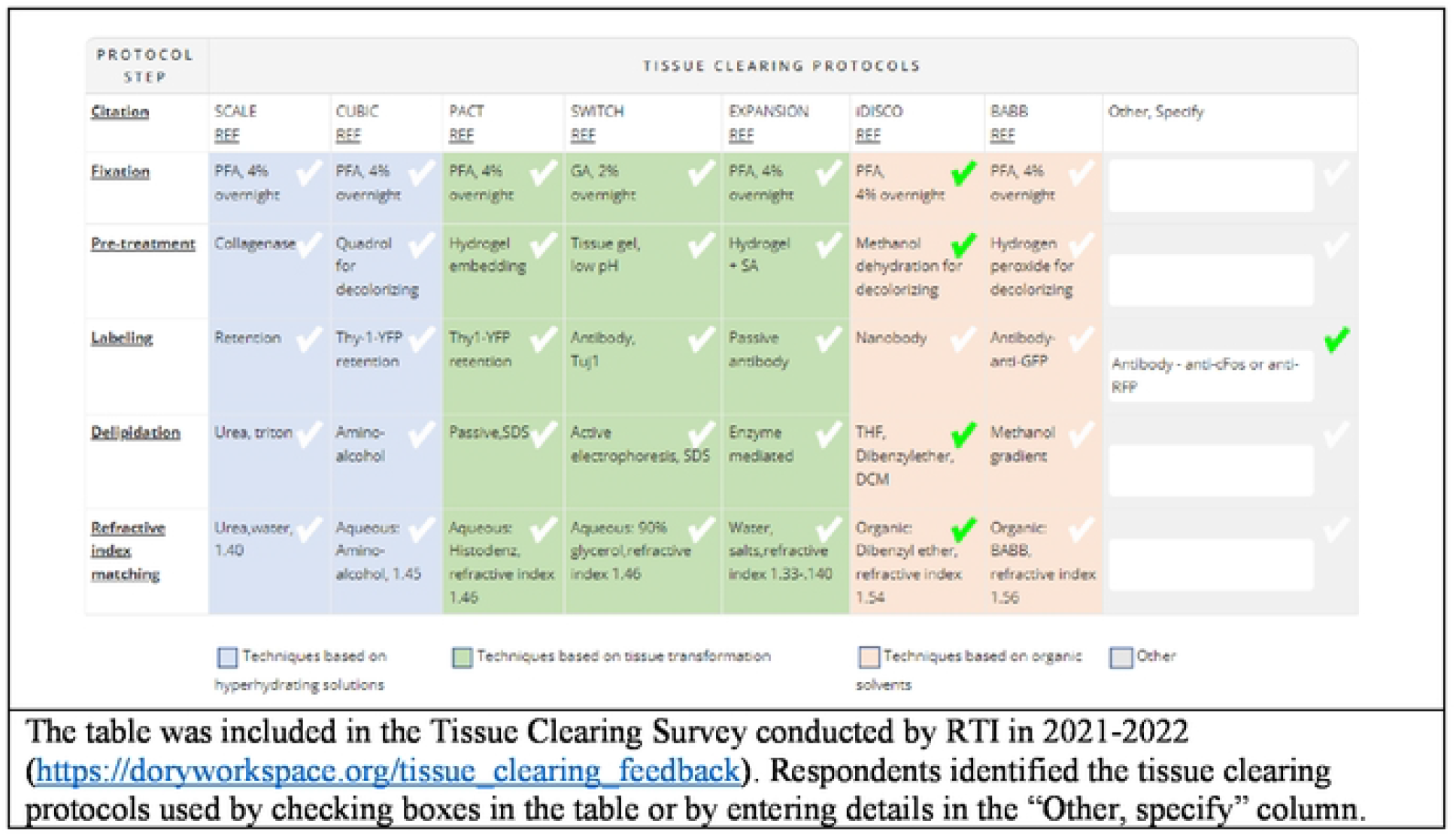
Table of Commonly Used Tissue-Clearing Protocols in the Tissue-Clearing Survey.

In an effort to increase response rate, a second outreach was performed between October 2021 and January 2022. Potential respondents were corresponding authors of publications identified by searching PubMed for keywords related to tissue clearing. The second outreach yielded 11 more responses.

### Analysis of Tissue-Clearing Survey Results

The results of the tissue-clearing survey were downloaded from the DORy website in CSV format and imported into Microsoft Excel®. Each unique entry describing a tissue-clearing protocol was recorded in a separate row. For each entry, the response for each survey question is recorded in a separate column (see complete dataset here: https://doryworkspace.org/doryviz).

Each unique tissue-clearing protocol was diagramed as a flow chart using Lucidchart software (**Figure 2**). Each flow chart included reported details for five major tissue-clearing steps) and the organism and tissue being cleared. Each protocol flow chart was organized according to whether the respondent indicated the protocol was very successful, successful, or not successful.

**Fig 2.**
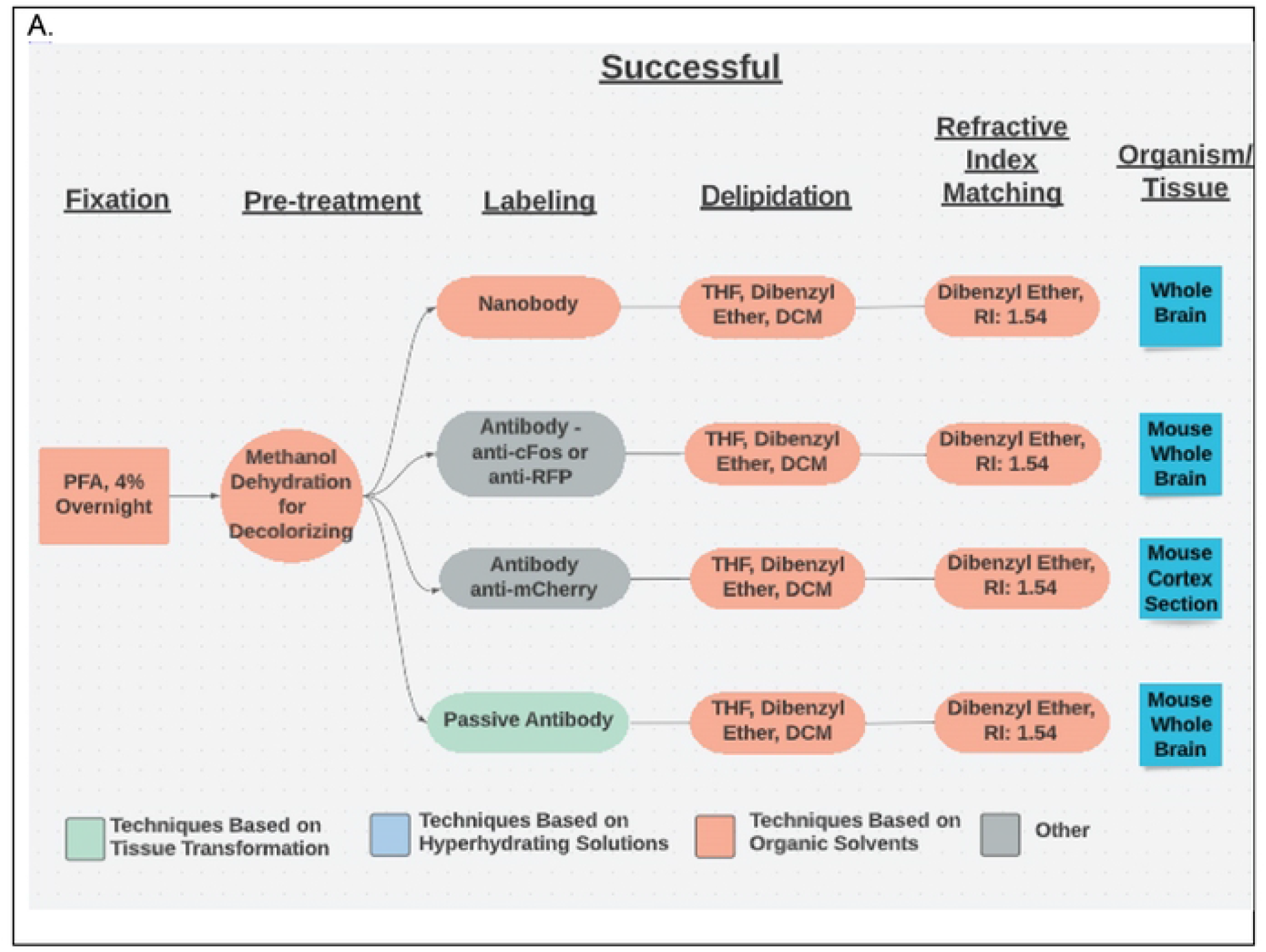
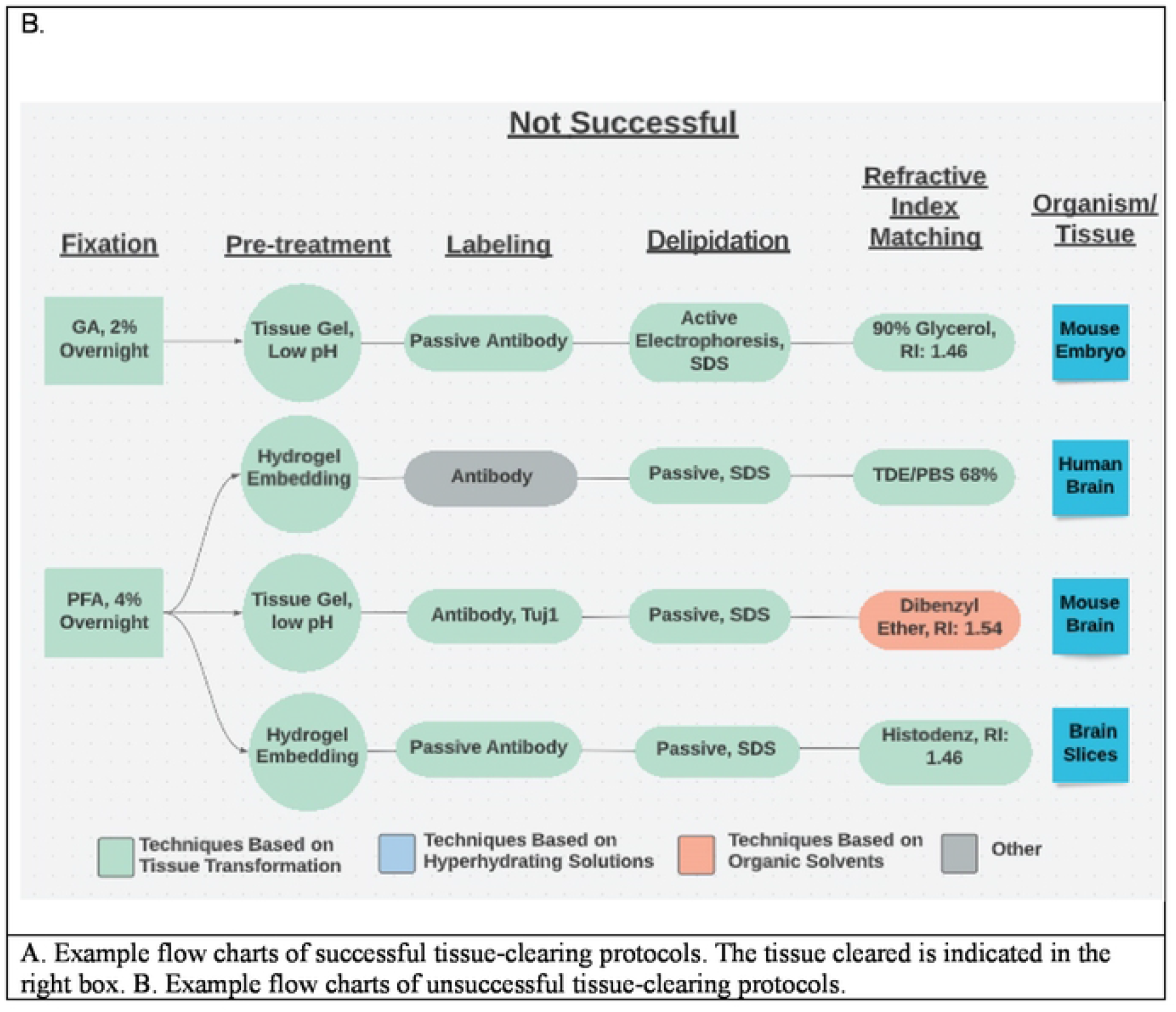
Example Flow Charts Created from Reported Tissue-Clearing Protocols.

### Development of T-CLEARE

Each flow chart was converted to a JavaScript Object Notation (JSON) tree data structure where the five different protocol steps were mapped to an arrays of key-value pairs (e.g., Fixation(*key*): PFA 4% overnight(*value*)). The JSON structure also contained some attributes to define the visualization part of the chart such as protocol step images, orientation of the nodes, or animation.

The individual JSON arrays were connected to a top-level parent “Protocol” node. The three children of the Protocol node are the key:value pairs indicating whether the protocols were “Very Successful,” “Successful,” and “Not Successful.” The fixation step is the child of protocol success, and each subsequent protocol step is the child of the previous protocol step (e.g., the delipidation step is the child of the fixation step). The tissue being cleared is the child of the refractive index matching protocol step (i.e., the leaf node).

The T-CLEARE presentation layer was implemented using pure JavaScript (JS) and the Treant.js (https://fperucic.github.io/treant-js/) open-source visualization library. The code included the jquery.min.js library to access and manipulate HTML elements. A custom JS library (dory-custom.js) was written to support the filters. CSS was used for styling. The T-CLEARE presentation layer was integrated into the Drupal-8 based DORy website (https://doryworkspace.org/doryviz) by packaging the JS and CSS files along with the JSON tree structure into a custom Drupal 8 module. T-CLEARE was tested with Chrome Version 101.0.4951.64 and is compatible with any browser (e.g., Firefox, Safari, and Microsoft Edge).

A standalone implementation of T-CLEARE is available from our GitHub repository (https://github.com/Defining-Our-Research-Methodology-DORy/TissueClearing). The lightweight T-CLEARE application can be run in any browser as a standalone client or integrated and deployed in other platforms.

## RESULTS

### T-CLEARE

T-CLEARE displays a tree diagram including each tissue-clearing protocol from the survey (**Figure 3**; https://doryworkspace.org/doryviz). When a user first opens T-CLEARE, the tree diagram displays four boxes. The “Protocols” box includes all flow diagrams for the reported tissue-clearing protocols. The next level to the right includes three boxes that organize the flow diagrams according to whether the protocols were “Very Successful,” “Successful,” or “Not Successful.” Clicking these three boxes will display the associated tissue-clearing protocol flow diagrams. The color of the box in the flow diagrams indicates whether the protocol step is associated with a hyperhydrating- (blue), tissue transformation- (aqua), or organic solvent-based class of tissue-clearing technique. Gray boxes indicate that the survey respondent used an “Other, specify” box to enter details for a protocol step. The fixation steps are yellow because these methods (e.g., PFA 4%, overnight) are shared among the classes of techniques.

**Fig 3.**
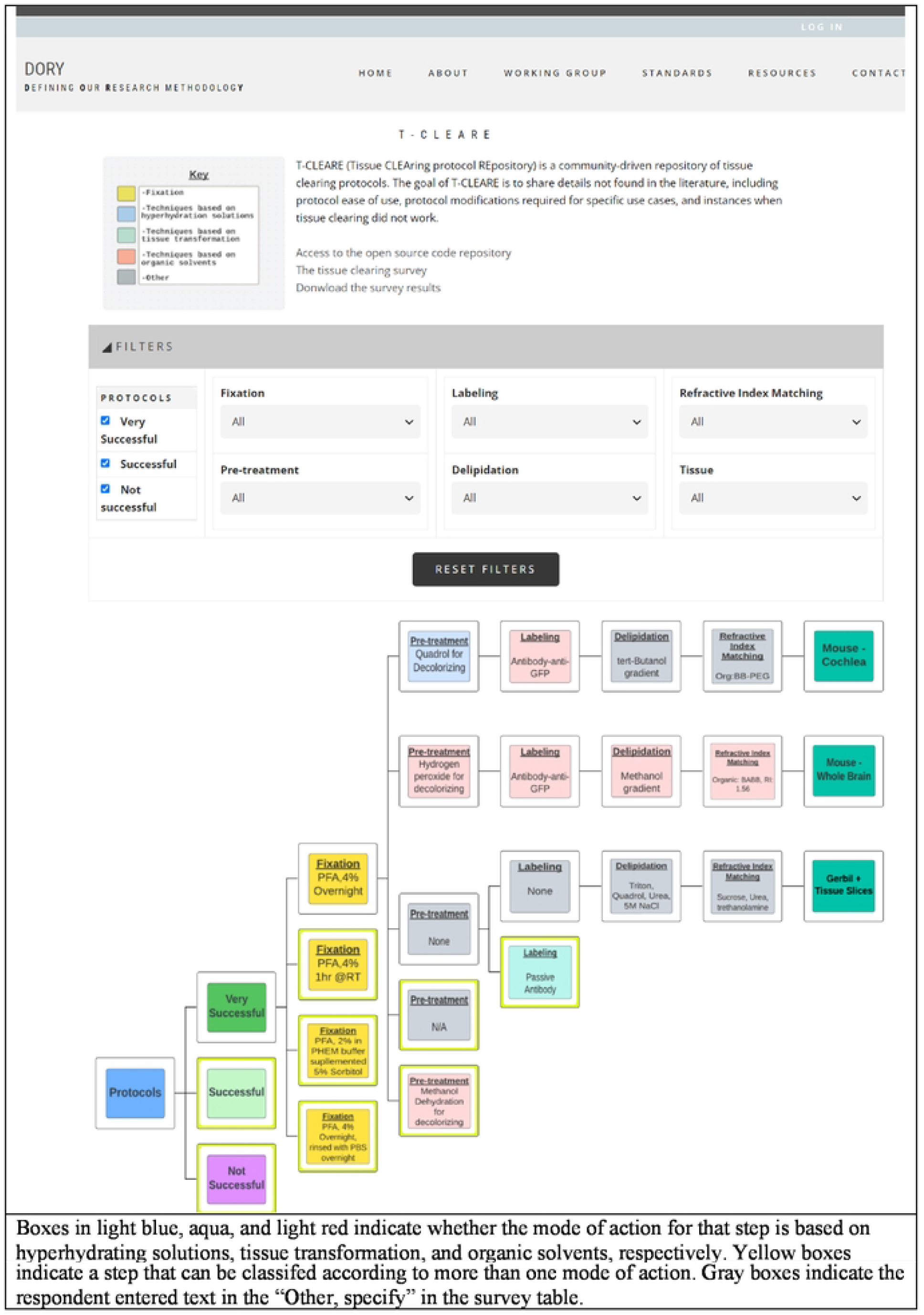
Screenshot of T-CLEARE User Interface.

The user can stratify the list of tissue-clearing flow diagrams displayed in the T-CLEARE using the filters at the top of the page. Available filters include level of success (e.g., very successful, successful, not successful), details for each of the five major steps of the protocol) and organism and tissue type (e.g., mouse–whole brain; mouse–cochlea). These filters can help users tease how successful a method or group of methods, is for a given tissue. For example, users can select “successful” protocols that follow hyperhydrating principles for fixation, pretreatment, labeling, and refractive index matching in “mouse brain.”

## DISCUSSION

Several limitations of the tissue-clearing survey results include a relatively small number of responses, the qualitative nature of the survey questions, overrepresentation of experiments in mouse brain (see **Figure 4**), and a large number of write-in responses, which are difficult to categorize. In spite of these limitations the results reflect a few overarching themes documented in the literature. The results support the assertion that most published tissue-clearing protocols require modification and optimization when applied to new experimental parameters (34). Of the 40 responses we received, 14 followed a tissue-clearing protocol as published (**Figure 5**). iDISCO (45) was most often cited as being followed without alterations (Figure 4), which supports observations that it is the most popular method reported in the literature (32). Twenty-seven either mixed steps from different published protocols or reported steps that differed from a published protocol (i.e., “other, specify”). This indicates that the optimal protocol for a given experiment may require blending reagents and steps from multiple established protocols (33, 34). Our survey results do not point to a single best technique that can be used across all experimental conditions (32, 46). **Figure 6** shows reported success of reported protocols when grouped according to mode of action (e.g., hyperhydrating techniques, tissue transformation techniques, organic solvent techniques). An examination of the 21 “successful” tissue-clearing protocols reveals relatively even distribution among hyperhydrating (six protocols), tissue transformation (eight protocols), and organic-solvent techniques (five protocols). On the other hand, 6 of the 11 tissue-clearing protocols that were “Very Successful” were based on organic solvent techniques while five of the eight protocols that were “Not Successful” were based on tissue transformation techniques. Because of the limited number of responses, it is difficult to determine if this indicates that techniques based on tissue transformation are not as effective as others or if this is just an artefact.

**Figure 4.**
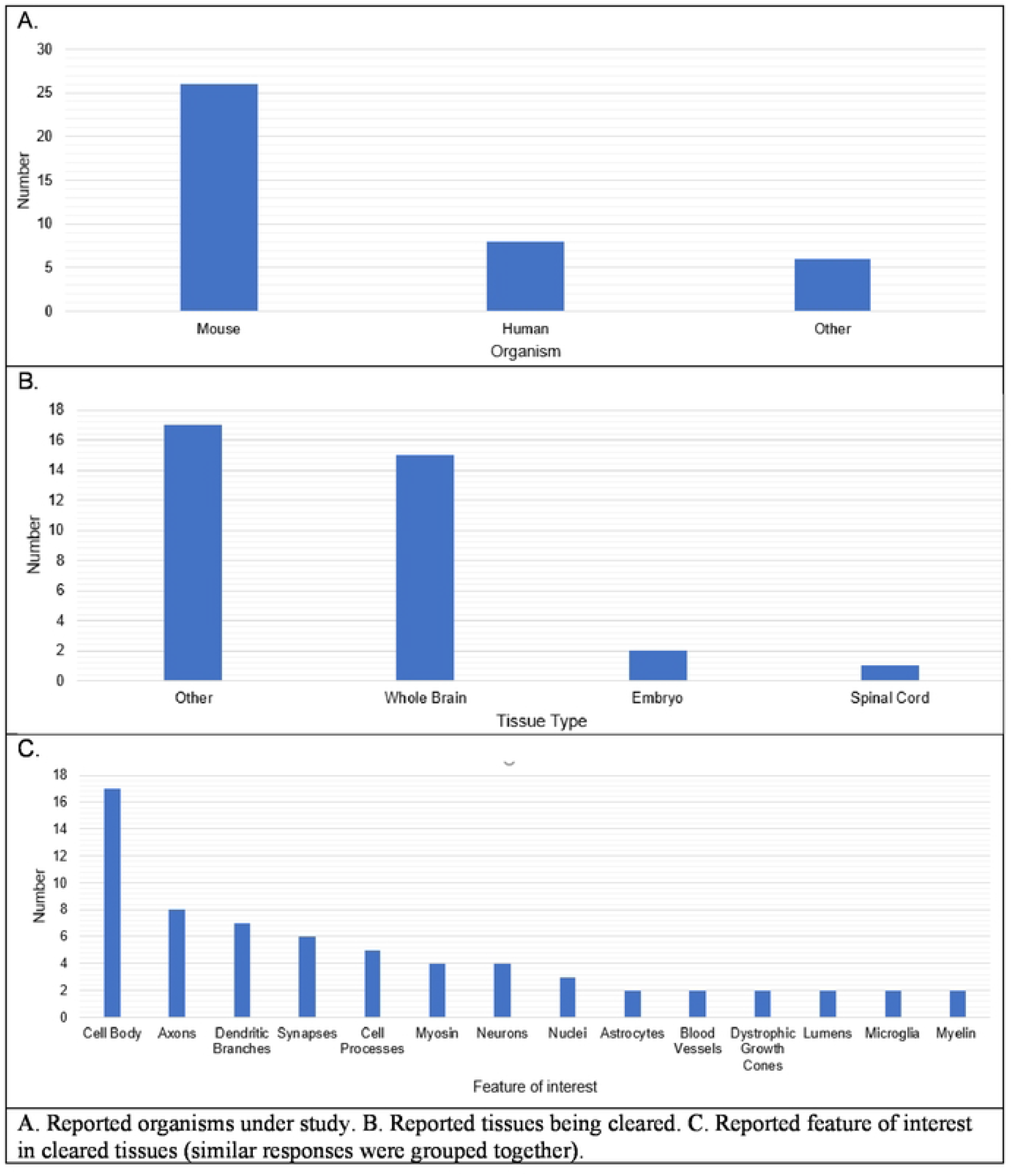
Organism, Tissue Type, and Feature of Interest in Reported Tissue Clearing Protocols.

**Fig 5.**
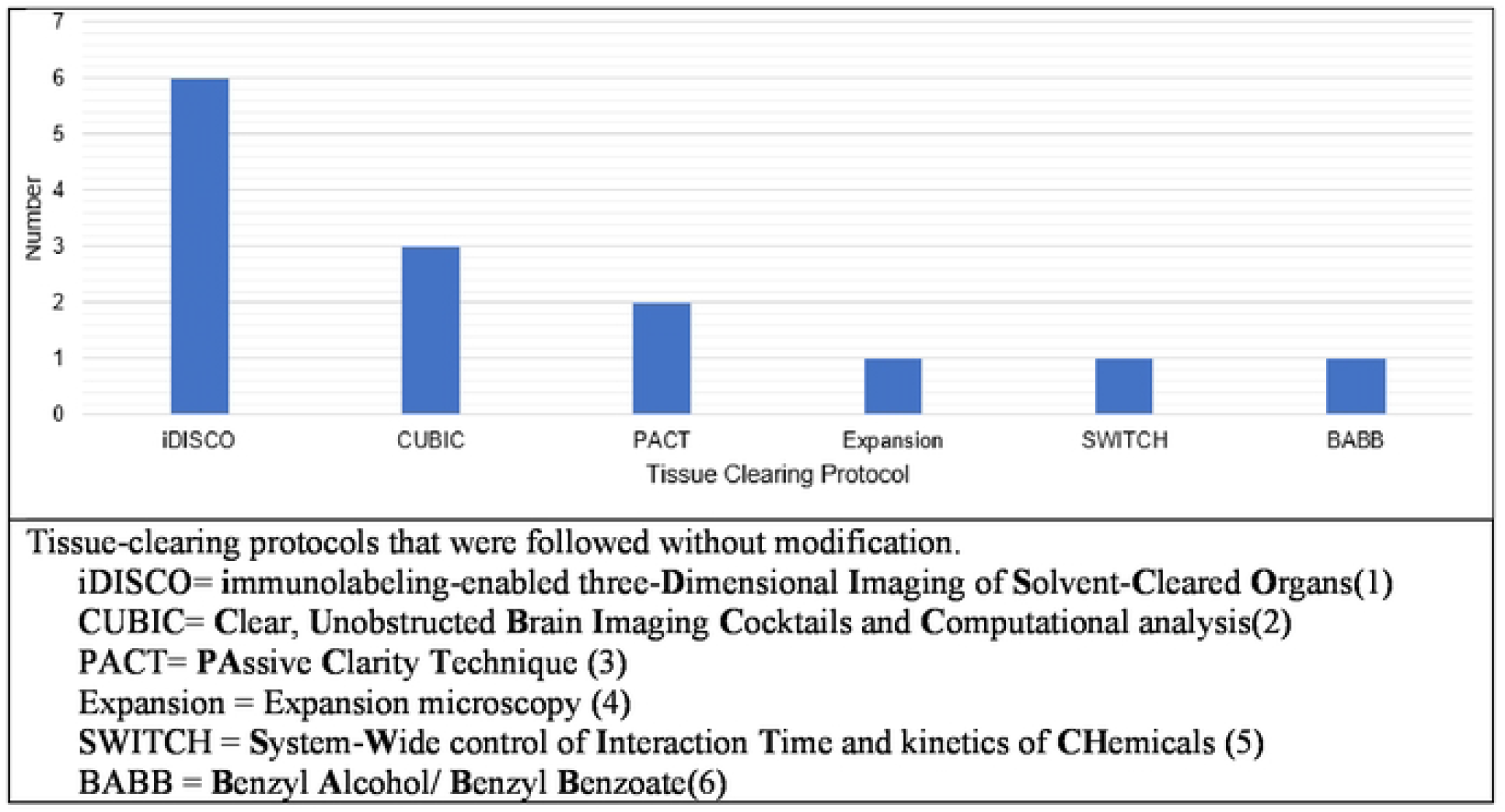
Reported Tissue-Clearing Protocols Used Without Modification.

**Fig 6.**
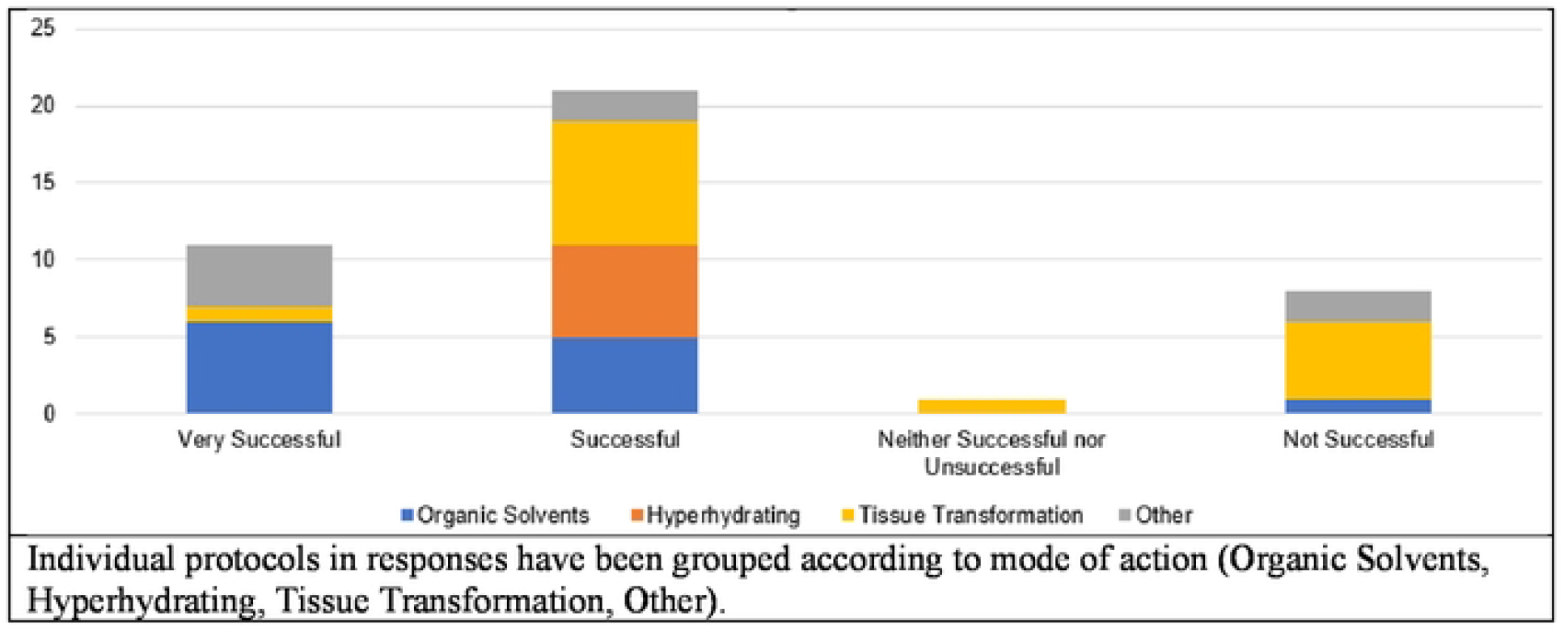
Overall Success in Reported Tissue-Clearing Protocols.

There are several ways to increase the utility of T-CLEARE. The most obvious is to gather additional input from the scientific community. To this end, the tissue-clearing survey will remain open and accessible from the DORy website (https://doryworkspace.org/tissue_clearing_feedback) for the community to report results of additional tissue-clearing experiments. There could also be future email campaigns asking for feedback on tissue clearing and other types of protocols (e.g., expansion microscopy). Future versions of T-CLEARE could automate the flow of the data from the survey to the user interface. Although the current lightweight JS/JSON implementation of T-CLEARE is modular (can be run standalone in any browser or easily integrated into other platforms) and scalable (additional protocols can be integrated as new JSON arrays), analyzing the survey results, curating the protocols flow diagrams, and creating images displayed in the nodes are manual steps. Future versions could include a MySQL database to capture and store the survey responses and algorithms that dynamically process new responses, generate the JSON arrays and images, and add them to the tree. Updates to the T-CLEARE user interface could include additional filters (e.g., microscope type used, type of analysis, level of analysis) and detailed comments from respondents. These comments, which provide details about protocol modifications, optimization, references, and why an experiment may have failed, are difficult to classify and are currently only available in the raw Excel data file. Future T-CLEARE features could link to full text versions of tissue-clearing protocols and resulting images in data repositories. The community is also encouraged to provide comments and suggestions using the feedback tools on the T-CLEARE and GitHUB repository pages.

In summary, T-CLEARE is a novel, web-based repository of tissue-clearing protocols to help the community share, find, and evaluate available tissue-clearing protocols in real time. As more information is added, we hope that T-CLEARE can help the community coalesce around reporting standards and standard tissue-clearing protocols. These community standards will help ensure consistent data collection and reporting, improve data interpretation, and facilitate data sharing among the scientific community.

## Acknowledgements

Research reported in this publication was supported by the National Institute of Mental Health (NIMH) of NIH under award number R24MH114683. The authors wish to acknowledge the members of the members of the BRAIN Initiative 3D Microscopy Working Group: Alex Ropelewski (co-chair), Hong-Wei Dong, Lydia Ng, Megan Rizzo, Jason Swedlow, Carol Thompson, and Pavel Osten. The authors also thank Neda Khanjani for technical expertise and Michelle Myers for expert manuscript editing. The content is solely the responsibility of the authors and does not necessarily represent the official views of NIH, NIMH, or any of the sponsoring organizations and agencies, or the U.S. government.

## Author Contributions

W.H. was the primary author on the manuscript. K.W. and J.H. led the development and refinement of the tissue-clearing survey, including the table of published tissue-clearing protocols. A.R., N.K., L.N., M.A.R., C.T., J.S., J.H., H.D., P.O., and K.W. are listed alphabetically and contributed to the manuscript either by their participation in the Working Group or by editing or commenting on the manuscript text. V.B., M.E., L.G., M.K., T.M., D.M., J.W., D.W., and C.H. contributed to the manuscript by supporting the Working Group consensus process or by editing and commenting on the manuscript.

